## Supplementary Data for "Leveraging place field repetition to understand positional versus nonpositional inputs to hippocampal field CA1"

**Supplementary Table 1 – Inclusion of Rewarded Alley Traversals Did Not Affect Main Results**

| Analysis, figure | Result excluding rewards (from main text) | Result including rewards |
|---|---|---|
| Proportion of fields with directional tuning (Fig 3B) | 93/310, binomial p = $3.6 \times 10^{-45}$ | 73/313, binomial p = $2.08 \times 10^{-28}$ |
| Lack of difference in directional tuning between repeating and nonrepeating neurons (Fig 4A) | Mann-Whitney U = 11883, p = 0.49 | Mann-Whitney U = 12067, p = 0.43 |
| Difference in correlation between directional tuning across fields in same versus different corridors (Fig 5E) | Real $r^2$ = 0.22,<br>$95^{th}$ percentile shuffle = 0.155 | Real $r^2$ = 0.26,<br>$95^{th}$ percentile shuffle = 0.156 |

Alley traversals on which the animal was rewarded were excluded from all main text analyses as any CA1 response to the reward itself was an undesirable source of variability. However, major analyses were re-run with rewards includeded (i.e. all passes through the alley) to test whether this changed the results. In each of three major analyses, the inclusion of reward trials did not qualitatively change the results compared to when these trials were excluded. It should be noted for the re-analysis of Fig 3B, the numerator decreased by a large amount despite little change to the overall pool of fields tested. The fact that including rewards did not result in many more fields being included can be explained by the fact that rewards were uniformly distributed across the track so their removal decreased the samples from all fields evenly, resuting in little change to the number of fields included for analysis. The fact that the numerator (i.e. directional fields) decreased could be explained by the fact that rewarded trials introduced variability, indeed this was the justificiation for their removal, and this variability reduced the statistical power of the Mann-Whitney test.

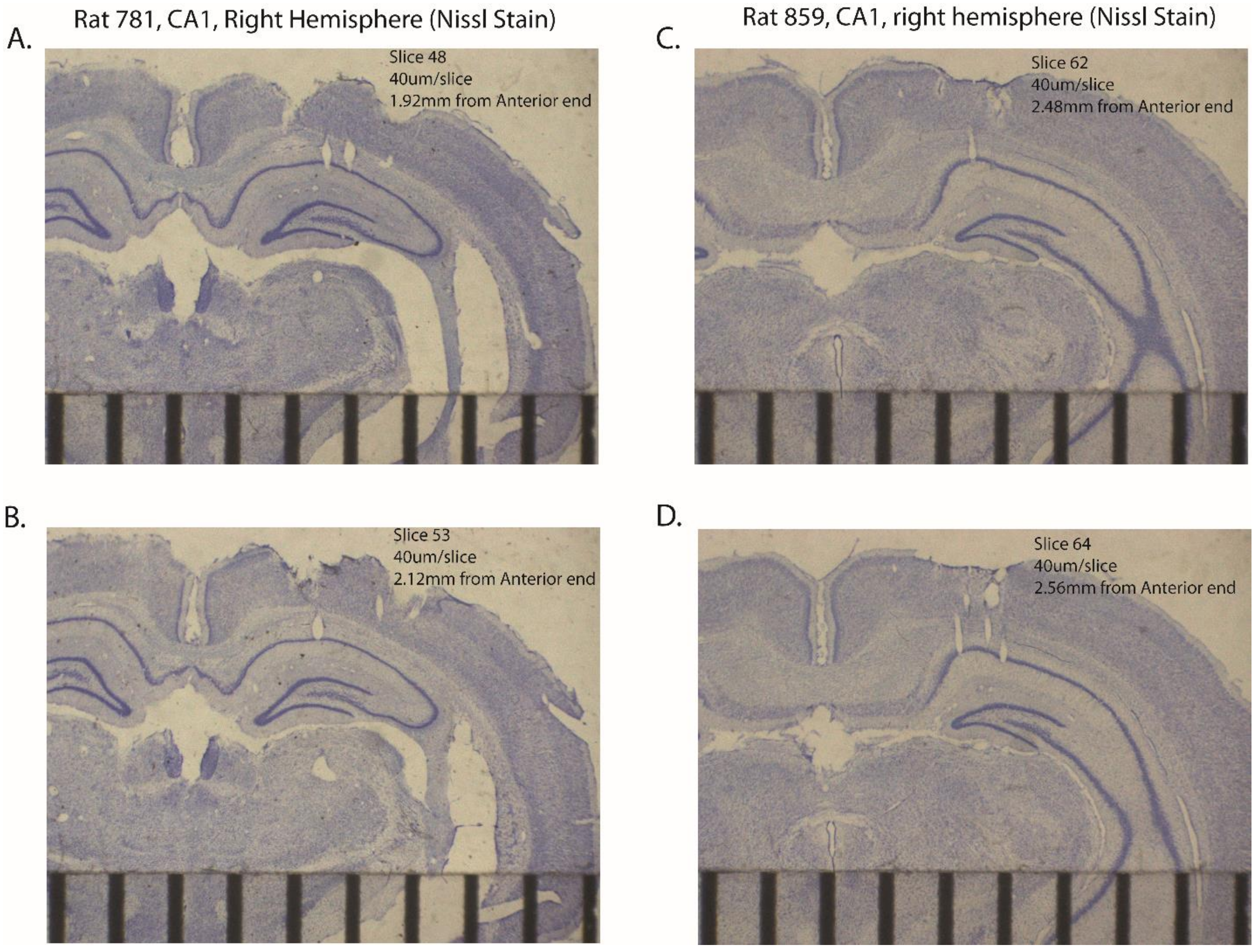


**Supplementary Figure 1 – Histology from Representative Animals.** All panels are 40 µm slices, Nissl stained from the right hemisphere CA1. Tears in/near CA1 layer indicate location of tetrode tracks. Ruler at bottom of figures shows mm gradations. **A.** Two tetrode tracks are visible in intermediate (along the transverse axis) CA1 at 1.92 mm along the anterior-posterior axis (A-P axis), starting from the septal pole of the hippocampus. **B.** One tetrode track visible in intermediate CA1 at 2.12 mm A-P. **C.** One tetrode track is visible in distal CA1 at 2.48 mm along the A-P axis. **D.** Three tetrode tracks are visible in/near distal CA1 at 2.56 mm A-P.

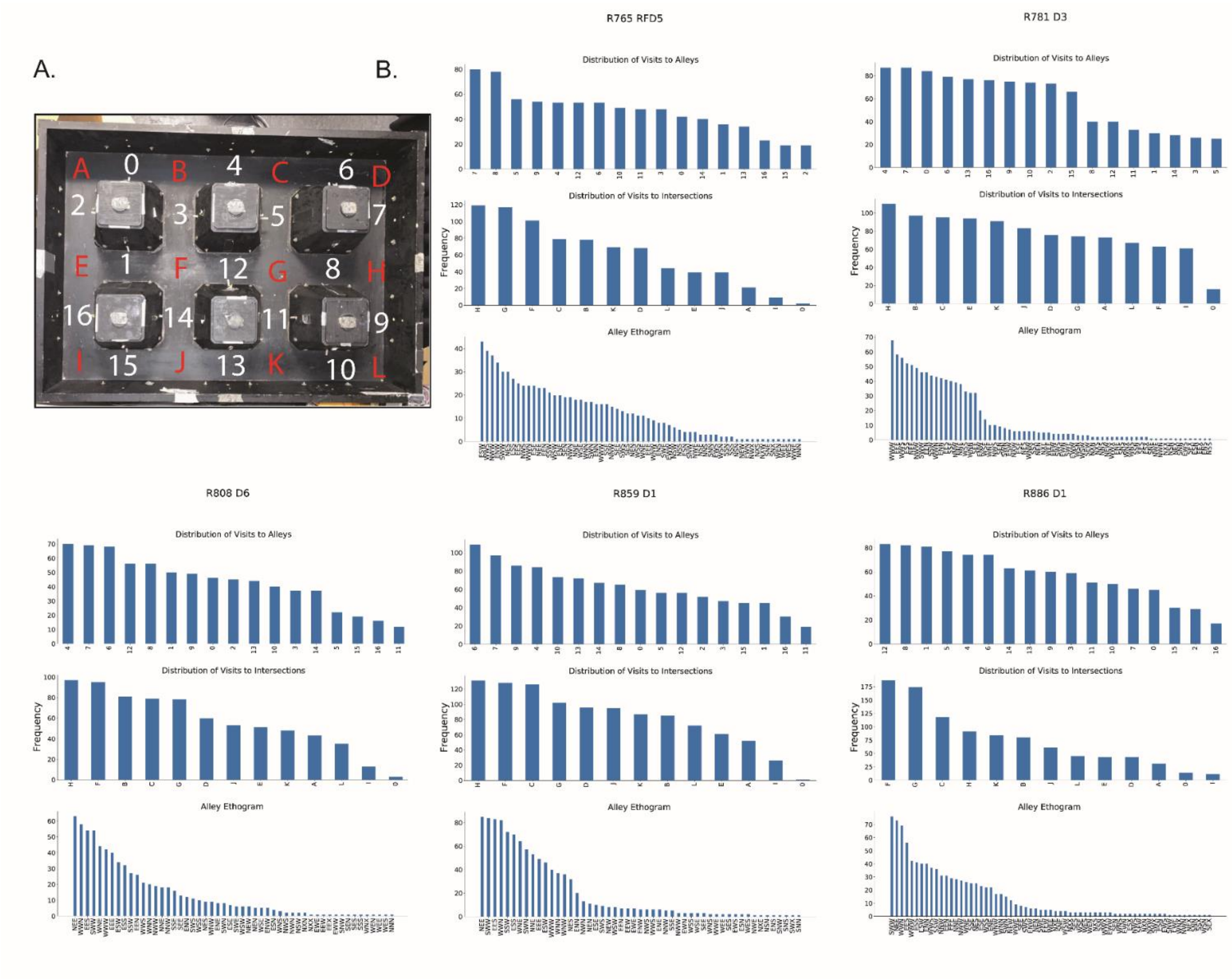


**Supplementary Figure 2 – Behavioral Sampling. A.** Photograph of city-block maze annotated with region labels. Alleys denoted by numbers, intersections by letters. **B.** Behavioral sampling of alleys and intersections for one day of recording of each rat. Top, histogram of visit counts to each alley. Middle, histogram of visit count to each intersection. Bottom, histogram of routes through alleys. A route is defined as a sequence of headings through a field: previous direction, current direction, next direction. For example, a label “NES” corresponds to a journey through the alley where the direction prior to turning into the alley was North, the path through the alley was East, and the turn out of the alley was South.

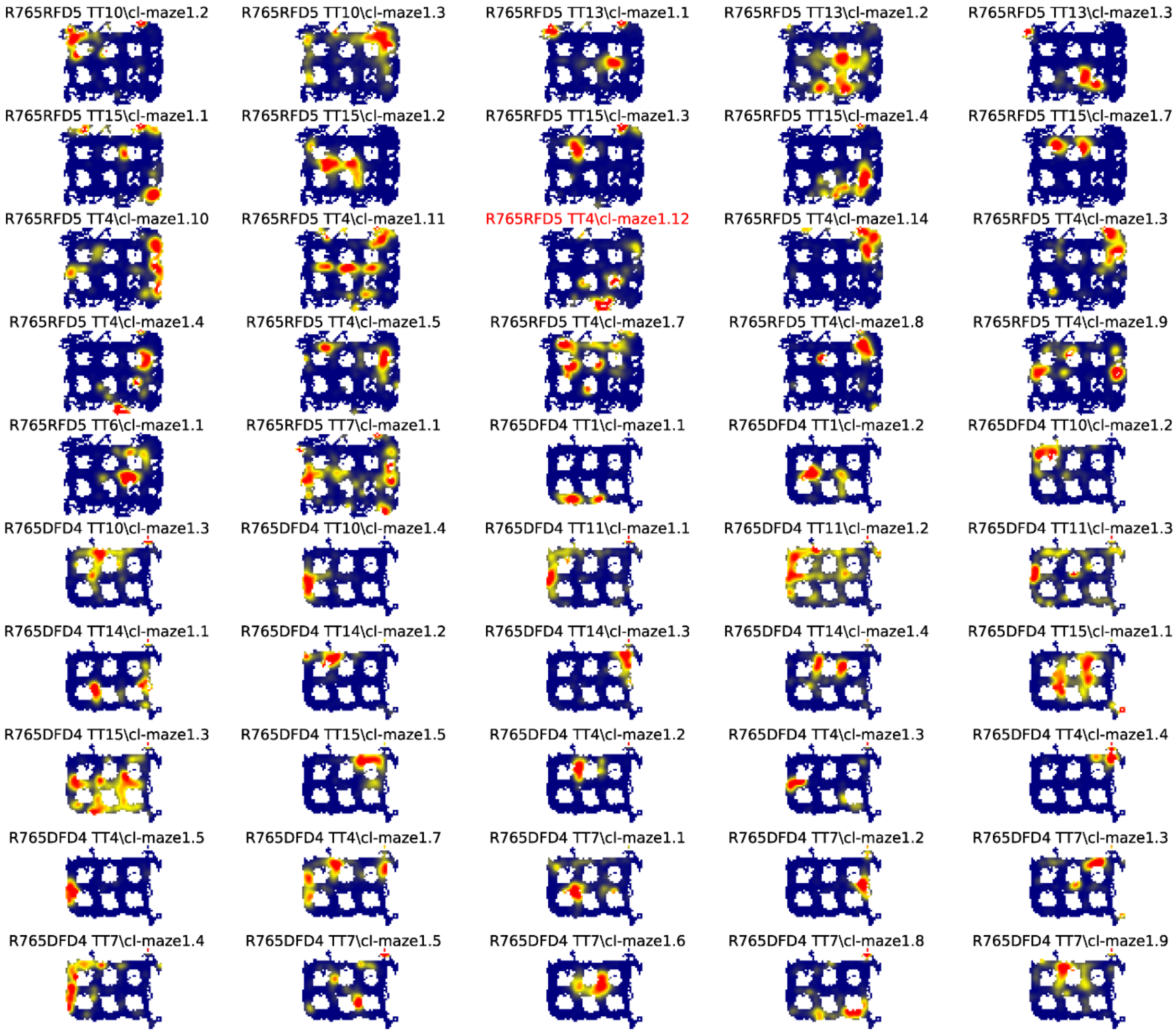
R765RFD5 TT10\cl-maze1.2
R765RFD5 TT10\cl-maze1.3
R765RFD5 TT13\cl-maze1.1
R765RFD5 TT13\cl-maze1.2
R765RFD5 TT13\cl-maze1.3
R765RFD5 TT15\cl-maze1.1
R765RFD5 TT15\cl-maze1.2
R765RFD5 TT15\cl-maze1.3
R765RFD5 TT15\cl-maze1.4
R765RFD5 TT15\cl-maze1.7
R765RFD5 TT4\cl-maze1.10
R765RFD5 TT4\cl-maze1.11
R765RFD5 TT4\cl-maze1.12
R765RFD5 TT4\cl-maze1.14
R765RFD5 TT4\cl-maze1.3
R765RFD5 TT4\cl-maze1.4
R765RFD5 TT4\cl-maze1.5
R765RFD5 TT4\cl-maze1.7
R765RFD5 TT4\cl-maze1.8
R765RFD5 TT4\cl-maze1.9
R765RFD5 TT6\cl-maze1.1
R765RFD5 TT7\cl-maze1.1
R765DFD4 TT1\cl-maze1.1
R765DFD4 TT1\cl-maze1.2
R765DFD4 TT10\cl-maze1.2
R765DFD4 TT10\cl-maze1.3
R765DFD4 TT10\cl-maze1.4
R765DFD4 TT11\cl-maze1.1
R765DFD4 TT11\cl-maze1.2
R765DFD4 TT11\cl-maze1.3
R765DFD4 TT14\cl-maze1.1
R765DFD4 TT14\cl-maze1.2
R765DFD4 TT14\cl-maze1.3
R765DFD4 TT14\cl-maze1.4
R765DFD4 TT15\cl-maze1.1
R765DFD4 TT15\cl-maze1.3
R765DFD4 TT15\cl-maze1.5
R765DFD4 TT4\cl-maze1.2
R765DFD4 TT4\cl-maze1.3
R765DFD4 TT4\cl-maze1.4
R765DFD4 TT4\cl-maze1.5
R765DFD4 TT4\cl-maze1.7
R765DFD4 TT7\cl-maze1.1
R765DFD4 TT7\cl-maze1.2
R765DFD4 TT7\cl-maze1.3
R765DFD4 TT7\cl-maze1.4
R765DFD4 TT7\cl-maze1.5
R765DFD4 TT7\cl-maze1.6
R765DFD4 TT7\cl-maze1.8
R765DFD4 TT7\cl-maze1.9

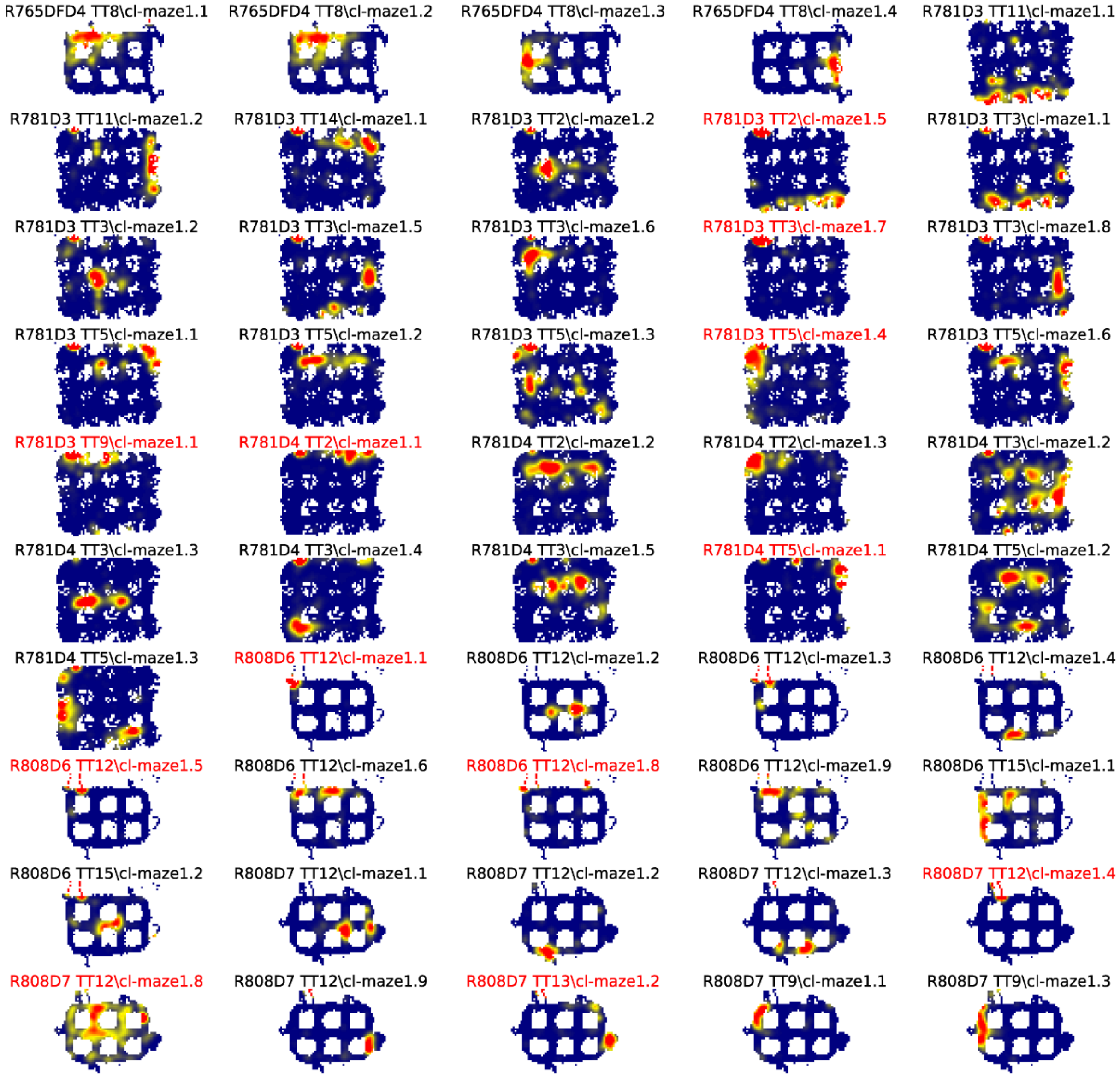
R765DFD4 TT8\cl-maze1.1
R765DFD4 TT8\cl-maze1.2
R765DFD4 TT8\cl-maze1.3
R765DFD4 TT8\cl-maze1.4
R781D3 TT11\cl-maze1.1
R781D3 TT11\cl-maze1.2
R781D3 TT14\cl-maze1.1
R781D3 TT2\cl-maze1.2
R781D3 TT2\cl-maze1.5
R781D3 TT3\cl-maze1.1
R781D3 TT3\cl-maze1.2
R781D3 TT3\cl-maze1.5
R781D3 TT3\cl-maze1.6
R781D3 TT3\cl-maze1.7
R781D3 TT3\cl-maze1.8
R781D3 TT5\cl-maze1.1
R781D3 TT5\cl-maze1.2
R781D3 TT5\cl-maze1.3
R781D3 TT5\cl-maze1.4
R781D3 TT5\cl-maze1.6
R781D3 TT9\cl-maze1.1
R781D4 TT2\cl-maze1.1
R781D4 TT2\cl-maze1.2
R781D4 TT2\cl-maze1.3
R781D4 TT3\cl-maze1.2
R781D4 TT3\cl-maze1.3
R781D4 TT3\cl-maze1.4
R781D4 TT3\cl-maze1.5
R781D4 TT5\cl-maze1.1
R781D4 TT5\cl-maze1.2
R781D4 TT5\cl-maze1.3
R808D6 TT12\cl-maze1.1
R808D6 TT12\cl-maze1.2
R808D6 TT12\cl-maze1.3
R808D6 TT12\cl-maze1.4
R808D6 TT12\cl-maze1.5
R808D6 TT12\cl-maze1.6
R808D6 TT12\cl-maze1.8
R808D6 TT12\cl-maze1.9
R808D6 TT15\cl-maze1.1
R808D6 TT15\cl-maze1.2
R808D7 TT12\cl-maze1.1
R808D7 TT12\cl-maze1.2
R808D7 TT12\cl-maze1.3
R808D7 TT12\cl-maze1.4
R808D7 TT12\cl-maze1.8
R808D7 TT12\cl-maze1.9
R808D7 TT13\cl-maze1.2
R808D7 TT9\cl-maze1.1
R808D7 TT9\cl-maze1.3

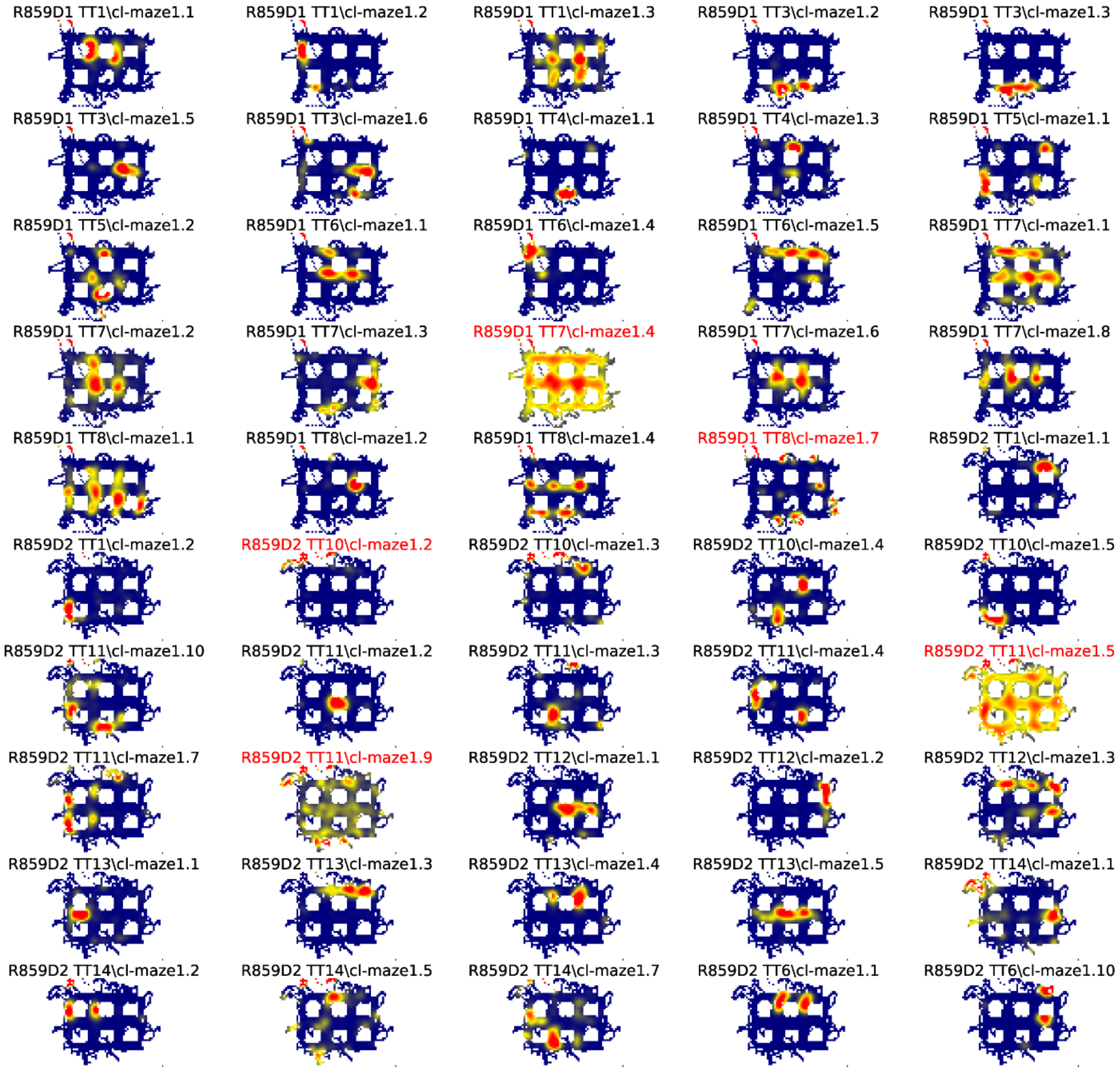
R859D1 TT1\cl-maze1.1
R859D1 TT1\cl-maze1.2
R859D1 TT1\cl-maze1.3
R859D1 TT3\cl-maze1.2
R859D1 TT3\cl-maze1.3
R859D1 TT3\cl-maze1.5
R859D1 TT3\cl-maze1.6
R859D1 TT4\cl-maze1.1
R859D1 TT4\cl-maze1.3
R859D1 TT5\cl-maze1.1
R859D1 TT5\cl-maze1.2
R859D1 TT6\cl-maze1.1
R859D1 TT6\cl-maze1.4
R859D1 TT6\cl-maze1.5
R859D1 TT7\cl-maze1.1
R859D1 TT7\cl-maze1.2
R859D1 TT7\cl-maze1.3
R859D1 TT7\cl-maze1.4
R859D1 TT7\cl-maze1.6
R859D1 TT7\cl-maze1.8
R859D1 TT8\cl-maze1.1
R859D1 TT8\cl-maze1.2
R859D1 TT8\cl-maze1.4
R859D1 TT8\cl-maze1.7
R859D2 TT1\cl-maze1.1
R859D2 TT1\cl-maze1.2
R859D2 TT10\cl-maze1.2
R859D2 TT10\cl-maze1.3
R859D2 TT10\cl-maze1.4
R859D2 TT10\cl-maze1.5
R859D2 TT11\cl-maze1.10
R859D2 TT11\cl-maze1.2
R859D2 TT11\cl-maze1.3
R859D2 TT11\cl-maze1.4
R859D2 TT11\cl-maze1.5
R859D2 TT11\cl-maze1.7
R859D2 TT11\cl-maze1.9
R859D2 TT12\cl-maze1.1
R859D2 TT12\cl-maze1.2
R859D2 TT12\cl-maze1.3
R859D2 TT13\cl-maze1.1
R859D2 TT13\cl-maze1.3
R859D2 TT13\cl-maze1.4
R859D2 TT13\cl-maze1.5
R859D2 TT14\cl-maze1.1
R859D2 TT14\cl-maze1.2
R859D2 TT14\cl-maze1.5
R859D2 TT14\cl-maze1.7
R859D2 TT6\cl-maze1.1
R859D2 TT6\cl-maze1.10

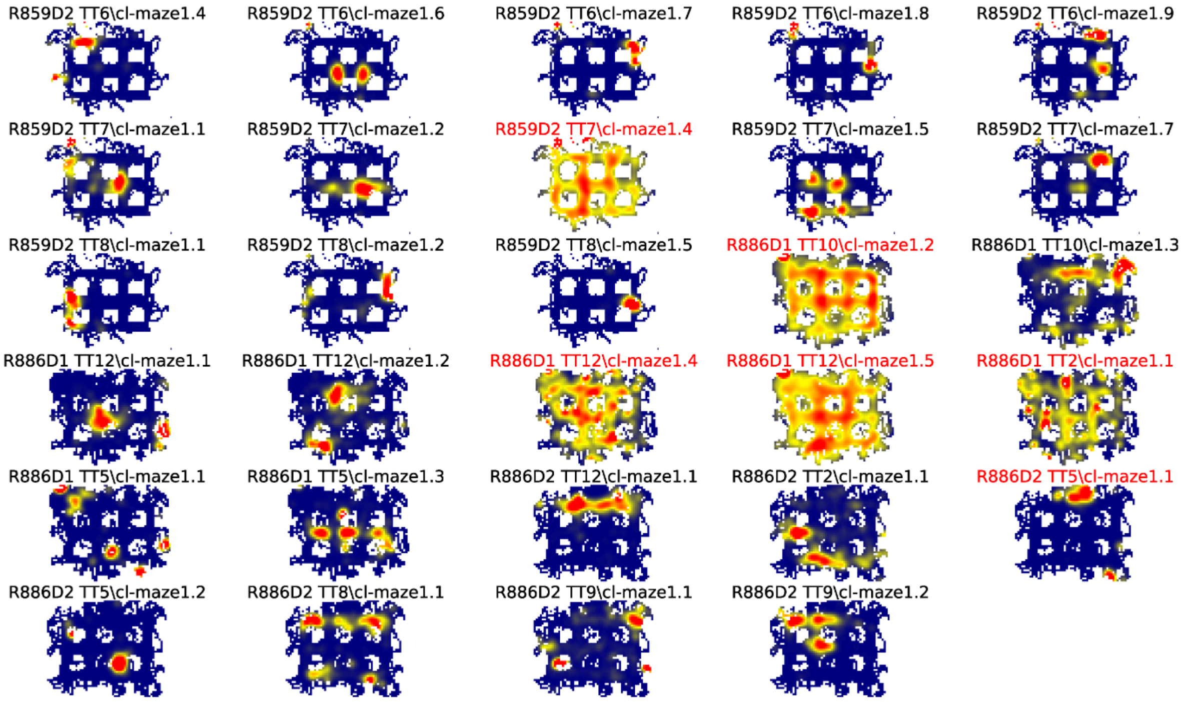


**Supplementary Figure 3 – All Ratemaps.** Each panel is the ratemap from a well-isolated neuron with on-track firing. No pass-sampling thresholds have been applied. Unlike the ratemaps shown in the main text, these ratemaps include the firing of the cell when the rat's head was outside the boundaries of the track, when it was rearing and investigating the outside environment rather than performing the task. The inclusion of this off-track behavior accounts for the ragged edges of the ratemaps. The off-track activity was not analyzed, due to the variability in this behavior across rats and across locations within rats. Instead, analysis was restricted to the alleys and intersections of the maze. Note the prevalence of cells that had multiple place fields in the alleys and intersections. Neurons excluded from futher analysis because they were classified as interneurons or did not have place fields are indicated by red titles.

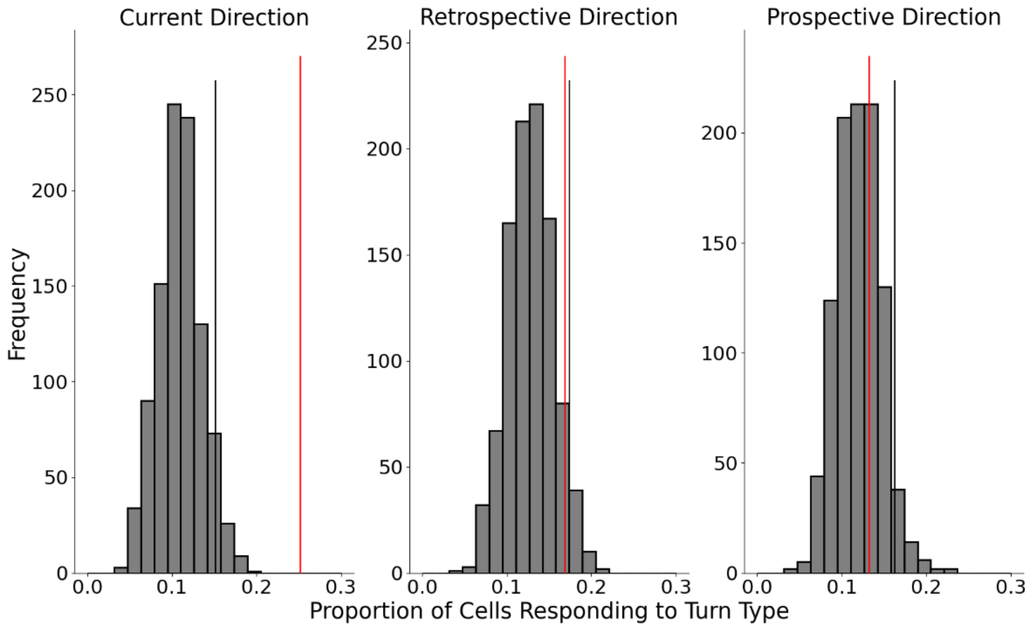


**Supplementary Figure 4 – Lack of Encoding of Retrospective or Prospective Direction**. A GLM approach was used to test for encoding of retrospective or prospective directions within single fields. The following model was fit to each field: Rate + 1 ~ $Dir_{Current}$ + $Dir_{Prospective}$ + $Dir_{Retrospective}$ + time, where time is represented as a natural spline with 3 df. The model is interpreted as testing the effect on firing rate of individual terms corresponding to previous, current, and next direction as well as time. The glm function from *stats* in R was used. Post-hoc comparison tests between levels of the same factor were performed using *emmeans* (Lenth et al., 2022). The proportion of fields with a significant effect of current direction (25.2%) was greater than the $95^{th}$ percentile of a shuffled distribution created by shuffling the sample firing rates from the sample direction labels. However, the proportions of fields with an effect of retrospective (16.9%) and prospective directions (13.2%) were not greater than the $95^{th}$ percentiles of their respective shuffled distributions. The effect of retrospective direction trended towards significance ($p = 0.069$).

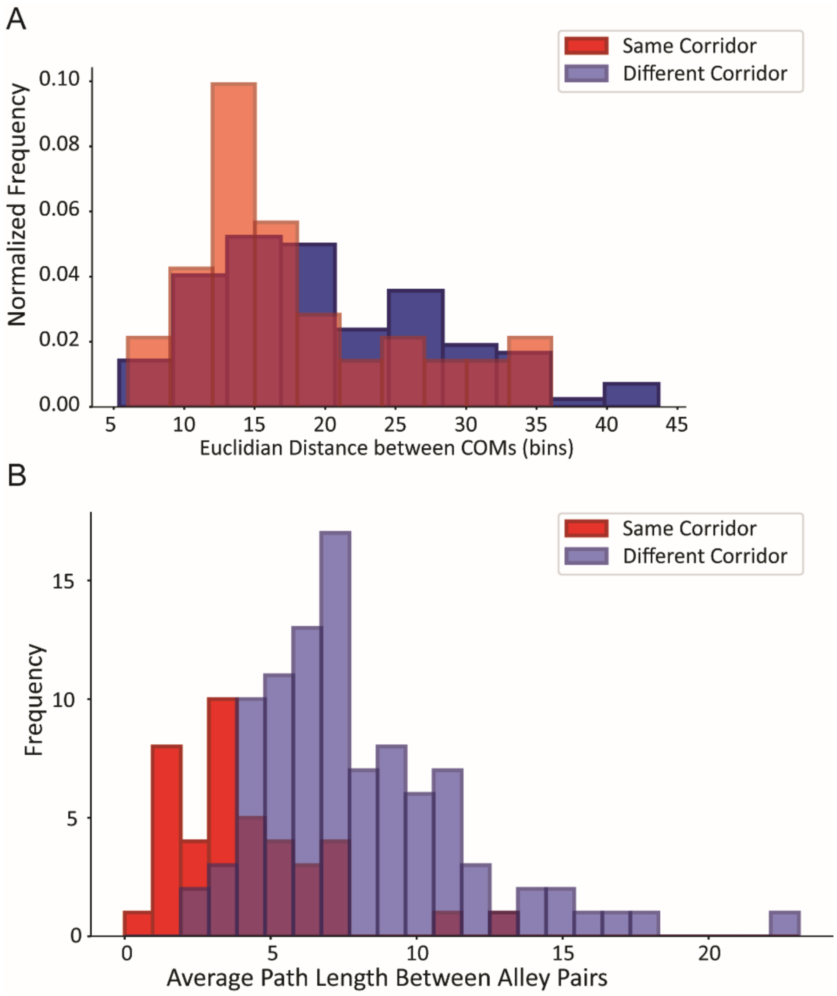


**Supplementary Figure 5 – Corridor Analysis Controls. A.** Distance between centers of mass (COMs) of fields comprising pairs on the same corridor (red) or different corridors (blue). Euclidian distance between field COMs (using straight-line distance between COMs, ignoring track geometry) on the same corridor was slightly shorter (17.55 ± 7.56 bins) than that of those on different corridors (20.1 ± 8.27 bins) (Mann Whitney U = 2097.5, p = 0.031) **B.** Travel distance between subsequent visits to fields comprising a pair on the same corridor (red) or a different corridor (blue). The unique pairs of alleys were used from each dataset based on all pairs of alleys on which repeating fields were located; that is, if 2 simultaneously recorded repeating cells had pairs of fields in the same alleys, that alley pair was used only once. For each alley pair, the average path length was calculated as the number of alleys separating successive visits to each alley of the pair, for all visits to the alleys. Each path length between alleys on different corridors was adjusted by -1 because it was impossible for there to be no intervening alleys for those paths (i.e. fields on alleys on different corridors could not be adjacent). The average path length for the same corridor pairs was shorter (4.17 ± 2.54 alleys) compared to different corridors (8.01 ± 3.47 alleys) (Mann-Whitney U = 607, p = $8.04 \times 10^{-11}$).

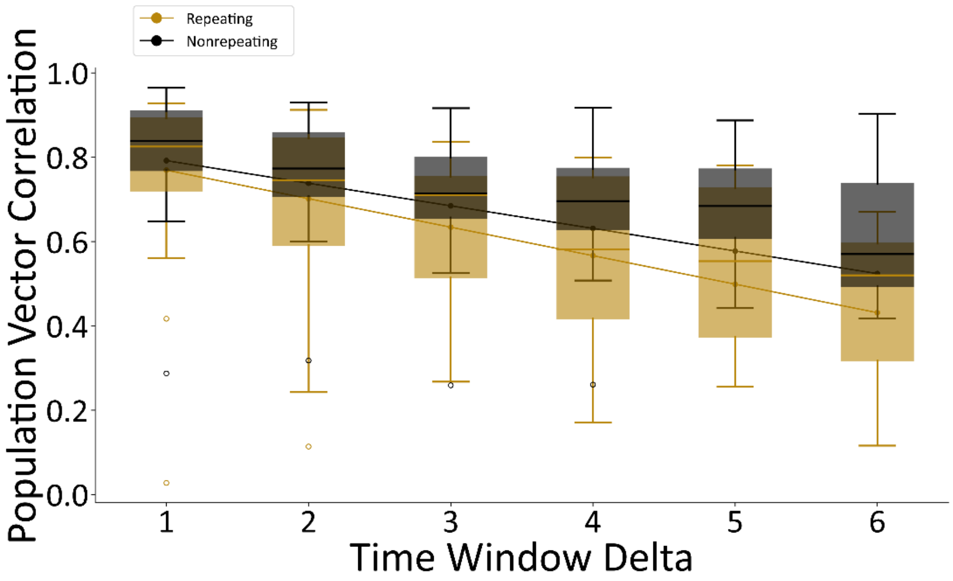


**Supplementary Figure 6 - Temporal Drift in Repeating vs Nonrepeating Neurons.** Population vector correlations as in Fig 6D, but done separately for repeating and nonrepeating fields. Linear regression model for repeating neurons: $r^2 = 0.20$, slope = -0.067, $p = 9.8 \times 10^{-8}$. Linear regression model for nonrepeating neurons: $r^2 = 0.16$, slope = -0.053, $p = 4.01 \times 10^{-12}$).

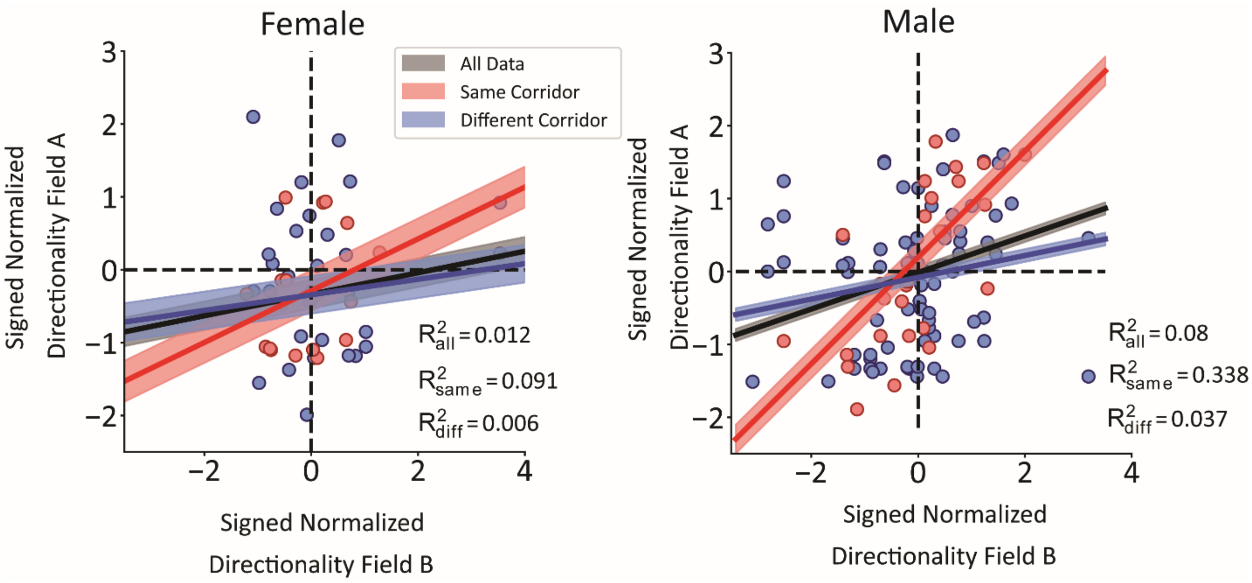


**Supplementary Figure 7 – Lack of Differences in Data within Male and Female Rats.** Rat sex was approximately balanced with 3 males and 2 females comprising the dataset. CA1 responses were similar within the main analyses presented here. Male and female rats did not have a statistically different number of operationally-defined repeating neurons (Male 26/97, female 30/82, Pearson's chi-square test $X^2(1) = 0.74$, $p = 0.38$). CA1 neurons were no more likely to be directionally tuned in male versus female rats (29.1% of male units (58/199) directional, 31.5% (35/111) of females, Pearson's chi-square test $X^2(1) = 0.039$, $p = 0.84$). Last, the figure panels show the results of the same corridor vs different corridor analysis. Although, the results were qualitatively similar between the sex groups, quantitative differences may be due to differences in sample size, and we do not place much confidence in this result based on data collected from only 2 female and 3 male rats.
